# NSun2 deficiency promotes tau hyperphosphorylation and neurodegeneration through epitranscriptomic regulation of miR-125b

**DOI:** 10.1101/2021.06.16.448614

**Authors:** Yoon A. Kim, Jennifer Blaze, Tristan Winters, Atul Kumar, Ellen Tein, Andrew A. Sproul, Andrew F. Teich, Francesca Bartolini, Schahram Akbarian, Gunnar Hargus, Ismael Santa-Maria

**Affiliations:** Taub Institute for Research on Alzheimer’s Disease & the Aging Brain, Columbia University, New York, NY, USA; Department of Pathology & Cell Biology, Columbia University, New York, NY, USA; Friedman Brain Institute, Icahn School of Medicine at Mount Sinai, New York, NY, USA; Department of Psychiatry, Icahn School of Medicine at Mount Sinai, New York, NY, USA

**Keywords:** Alzheimer’s disease, NSun2, neurodegeneration, Tau proteostasis, microRNA, methylation

## Abstract

Overproduction or suppression of certain microRNAs (miRNAs) in Alzheimer’s disease (AD) brains promote alterations in tau proteostasis and neurodegeneration. However, the mechanisms governing how specific miRNAs are dysregulated in AD brains are still under investigation. Epitranscriptomic regulation adds a layer of post-transcriptional control to brain function during development and adulthood. NOP2/Sun RNA methyltransferase 2 (NSun2) is one of the few known brain-enriched methyltransferases able to modify mammalian non-coding RNAs and loss of function autosomal-recessive mutations in NSUN2 have been associated with neurological abnormalities in humans. Here, we provide evidence that NSun2 is expressed in adult human neurons in the hippocampal formation and prefrontal cortex. When we evaluated NSun2 protein expression in *post-mortem* brain tissue from AD patients we find is dysregulated which was also found in mice and human cellular AD models. To probe these observed alterations were unique to AD we further evaluated brain tissue from other tauopathies, observing NSun2 protein levels were similar between cases and controls. In a well-established *Drosophila* melanogaster model of tau-induced toxicity we investigated the pathological role of NSun2 observing that reduction of NSun2 protein levels exacerbated tau toxicity, while overexpression of NSun2 partially abrogated the toxic effects. We further show using human induced pluripotent stem cell (iPSC) derived neuronal cultures that NSun2 deficiency results in tau hyperphosphorylation and we found in primary hippocampal neuronal cultures NSun2 levels decrease in response to amyloid-beta oligomers (AβO). Furthermore, in mice, we observed that NSun2 deficiency promotes aberrant levels of m6A methylated miR-125b and tau hyperphosphorylation. Altogether, our study supports that neuronal NSun2 deficiency in AD promotes neurodegeneration by altering tau phosphorylation and tau toxicity through an epitranscriptomic regulatory mechanism and highlights a novel avenue for therapeutic targeting.

## Introduction

MiRNAs, a class of non-coding small RNAs, are part of a vital regulatory mechanism that prevents the deposition of tau protein. Several miRNAs, have been shown to regulate tau proteostasis by modulating tau synthesis or post-translational modifications on tau, such as phosphorylation^1, 2^. However, mechanisms governing how miRNAs are regulated in the brain or how they are dysregulated during the disease process are poorly understood^3–12^. MiRNAs can be regulated at the transcriptional or post-transcriptional level^13, 14^ One of the most frequent post-transcriptional modifications of RNA is methylation^15–18^. However, the specific role that RNA methyltransferases play in neurodegeneration is poorly investigated. NSun2 is one of the few brain-enriched methyltransferases known to facilitate methylation of non-coding RNAs, including microRNAs^19–23^. Based on the previously reported putative neuroprotective role of NSun2 and its constitutive expression in the mouse brain cortex and hippocampus^19, 24, 25^, we decided to explore the status of NSun2 in AD models and human tissue. Here, we show that Nsun2 is downregulated in AD, modulates tau toxicity *in vivo*, and regulates tau phosphorylation in part by promoting epitranscriptomic alterations in miR-125b.

## Results

### NSun2 RNA methyltransferase is dysregulated in Alzheimer’s Disease brains

*Post-mortem* examination of human control brains shows NSun2 positive immunostaining in neurons of the hippocampal formation and the prefrontal cortex (Brodmann area 9). Immunolabeling of the nucleus and dendrites shows for the first time that NSun2 protein is expressed in neurons in the adult human brain (**Supplementary Figure 1**). To determine whether alterations in NSun2 are found in AD we next performed NSun2 immunohistochemistry on brain sections from AD cases and controls (**Supplementary Table 1**). Prefrontal cortex and hippocampal formation were analyzed as these are some of the most affected brain regions in AD. The most salient feature was the neuronal nuclear decrease of NSun2 immunoreactivity in both brain regions of AD cases compared to controls (**Figure 1A**). Apart from the decrease of nuclear immunoreactivity, AD patients also show a decreased staining in the soma and neurites (right panels, **Figure 1A**).

**Figure 1.**
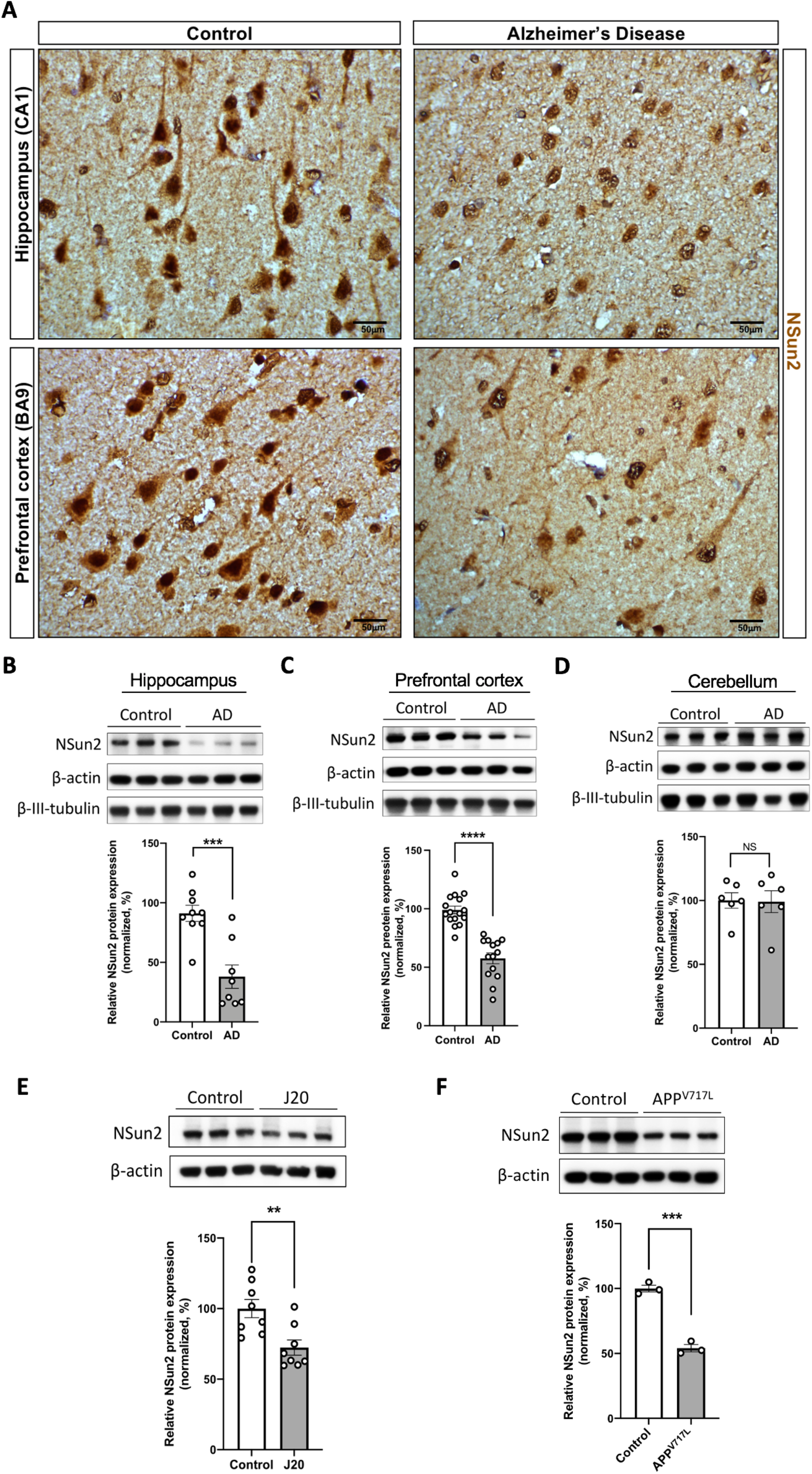
NSun2 RNA methyltransferase is dysregulated in Alzheimer’s Disease. (**A**) Representative NSun2 immunohistochemistry images of the hippocampal formation (Cornu Ammonis1 (CA1)) and prefrontal cortex (Brodmann area 9) of age-matched controls and AD human brains. Scale bars, 50 μm. (**B**) Western blot quantification of NSun2 protein levels in human hippocampus of controls (*n* = 9) and AD (*n* = 8). (**C**) Western blot quantification of NSun2 protein levels in the prefrontal cortex of AD patients (*n* = 14) compared to their respective controls (*n* = 16). (**D**) Western blot quantification of NSun2 protein levels in the cerebellum of controls (*n* = 6) and AD patients (*n* = 6). Mann Whitney *U* Test; \*\*\**P* < 0.001, \*\*\*\**P* < 0.0001. Data represent mean ± SEM. (**E**) Western blot quantification of NSun2 protein levels in the hippocampus of 6-month-old J20 mice (*n* = 8) and non-transgenic controls (*n* = 8). (**F**) Western blot quantitative analysis on APP^V717L^ (*n* = 3) and isogenic control (*n* = 3) iPSC-derived neurons for NSun2 protein. Student’s t test; \*\**P* < 0.01, \*\*\**P* < 0.001. Data represent mean ± SEM. (**B-F**) Top portion of the panels shows representative Western blots. (**B-G**) Histograms show densitometric quantification of NSun2 protein abundance with respect to control at the bottom of the panels. NSun2 is normalized by β-actin in all samples.

We then performed quantitative Western blot analysis to confirm whether levels of NSun2 are reduced in AD brains. Immunoblots using antisera specifically recognizing human NSun2^26, 27^ show a decrease in the levels of NSun2 both in hippocampal formation and the prefrontal cortex, in comparison to samples from control subjects (**Figure 1 B, C**). In this case, the reduction was more pronounced in the hippocampal formation (53.18% decrease, *P* = 0.0009) than in the prefrontal cortex (41.36% decrease, *P* <0.0001) (**Figure 1B, C**). Importantly, no significant difference was observed when comparing AD cases and controls in the cerebellum, a brain area devoid of AD pathology (**Figure 1D**). Notably, we did not detect a difference in the beta-III tubulin neuronal marker^28^ indicating that our observed differences might not be secondary to neuronal loss. However, AD brains did show significantly higher levels of NSun2 mRNA in the prefrontal cortex, although quantitative real-time PCR (QPCR) analysis did not show a significant difference in the levels of NSun2 mRNA in the hippocampus, suggesting a compensatory mechanism observable in the prefrontal cortex, a brain area that degenerates later in the disease process (**Supplementary Figure 2**). Furthermore, we analyzed publicly available proteomic and transcriptomic datasets^29, 30^ which showed results in agreement with our findings (**Supplementary Figure 3**).

It is unlikely that NSun2 alone among methyltransferases has the ability to regulate microRNAs. Indeed, a possible cooperative mechanism between NSun2 and Methyltransferase Like 3 (Mettl3) has been proposed^31^. To investigate whether AD brains also show biochemical changes in Mettl3 we examined the levels of Mettl3 expression in control and AD brains. Western blot analysis did not show a significant decrease in Mettl3 levels (**Supplementary Figure 4A, B**). In addition, analysis of a publicly available transcriptomic dataset did not show a significant change in Mettl3 mRNA levels corroborating our findings (**Supplementary Figure 4C**).

Next, we sought to confirm in two distinctive AD models whether NSun2 downregulation is also observed in these model systems. First, we analyzed the J20 mouse model of AD. This mouse model overexpresses human Amyloid Precursor Protein (APP) with two mutations (APP KM670/671NL-V717F) linked to familial AD^32^. J20 mice develop robust amyloid beta (Aβ) pathology by five to seven months of age, showing learning and memory deficits and changes in synaptic plasticity^32^. Thus, we performed Western blot analysis in hippocampal samples from 6-month-old J20 mice. We observed a significant reduction of NSun2 levels in the hippocampus compared to wild type controls (27.57% decrease, *P* = 0.0055) (**Figure 1E**). In addition, we were able to recapitulate the alterations in NSun2 observed in AD brains in induced pluripotent stem cell (iPSC) derived neurons from an AD heterozygous knockin hiPSC line (IMR90, cl.4 backbone, WiCell) harboring the APP V717L London mutation^33–39^. Neurons bearing this mutation show alterations in APP processing, and tau proteostasis^40, 41^. In this AD *in vitro* model, we observed a significant reduction in the levels on NSun2 when compared to isogenic controls (45.95% decrease, *P* = 0.0003) (**Figure 1F**).

To further explore whether downregulation of NSun2 is a common event among other tauopathies we performed immunohistochemistry and Western blot analysis on Primary Age-Related Tauopathy (PART)^42^ and Progressive Supranuclear Palsy (PSP) (**Supplementary Figure 5**)^43, 44^ using samples from the hippocampal formation and the Globus pallidus respectively; main areas affected in the brains of patients with these disorders^42, 44^. Quantitative Western blot analysis using NSun2 antisera showed no significant difference in the levels of NSun2 (**Supplementary Figure 5D, E**). These results suggest reduction of NSun2 protein levels in human brains is specific to AD when compared to other tauopathies.

### Deficiency of NSun2 promotes alterations in tau phosphorylation and tau toxicity

Next, we asked whether NSun2 influences tau toxicity *in vivo*. Many molecular mechanisms of post-transcriptional regulation including epitranscriptomic modifications and microRNA regulation are conserved between *Drosophila* melanogaster (fruit flies) and humans^45^. Furthermore, fruit flies have proven to be useful for modeling essential mechanisms of tauopathy and tau biology^46^. Using a conditional expression system, we overexpressed either NSun2 or a short interfering RNA (siRNA) against NSun2. First, human tau was co-expressed with a siRNA control in the *Drosophila* eye, producing a rough eye phenotype, confirming that the model system was functioning (**Figure 2A**). Next, human tau and NSun2 siRNA were co-expressed resulting in an exacerbation of the phenotype (**Figure 2A**). In contrast, coexpression of human tau and *Drosophila* NSun2 showed a partial reversal of the rough eye phenotype, consistent with a protective role, demonstrating bidirectionality (**Figure 2B**). Quantitative assessment of these phenotypes revealed that these findings are highly significant (**Figure 2A, B**), indicating that NSun2 influences tau toxicity in this system.

**Figure 2.**
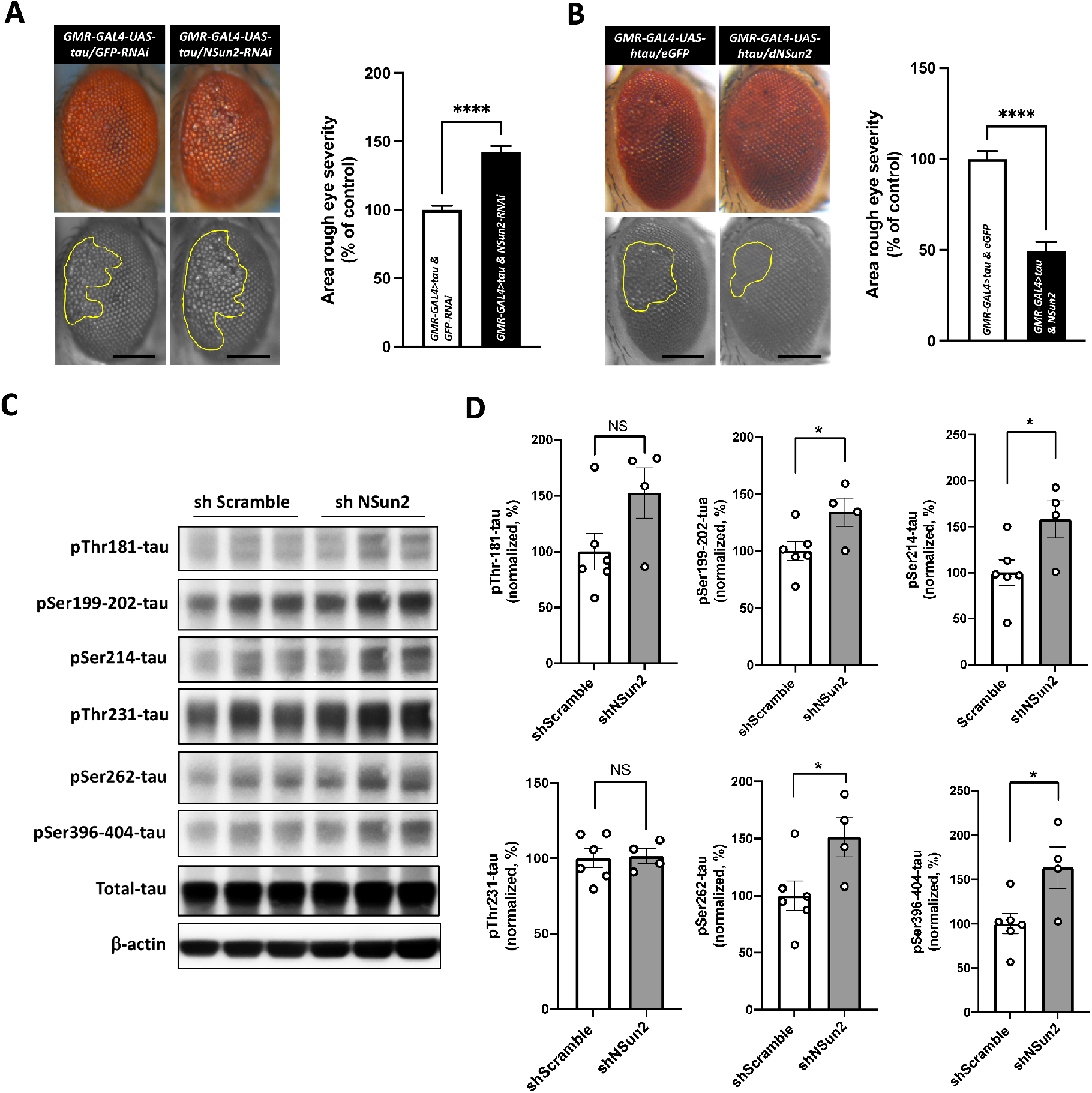
Regulation of tau toxicity and tau phosphorylation by NSun2. (**A**) Co-expression of NSun2 RNAi (*n* = 26) with human tau exacerbated the rough eye phenotype compared with that observed in the GFP RNAi control (*n* = 29). (**B**) Co-expression of NSun2 (*n* = 18) partially suppressed the human tau-induced rough eye phenotype compared with that seen in the GFP control (*n* = 28). The yellow marked area shows the degenerated part of eyes. Scale bars, 200 μm. Histograms show quantitative assessment of eye phenotypes (\*\*\*\**P* ≤ 0.0001 by Mann-Whitney *U* test). (**C**) human iPSC derived neurons were transduced with shNSun2 or scramble control, and protein lysates collected and analyzed. Representative Western blots with indicated antibodies demonstrated the effects of shNSun2 on the levels of phosphorylated forms of tau. (**D**) Quantification of phosphorylated tau in neurons transduced with shNSun2, plotted relative to the levels of total tau (after normalization of total tau with β-actin levels). (Student’s t test; \**P* < 0.05). Data represent mean ± SEM.

We next set out to investigate in human neurons whether reduction of NSun2 might modulate tau toxicity by altering tau phosphorylation levels. To this aim we took advantage of iPSC derived neurons as a favored *in vitro* model system to interrogate and investigate molecular events driving tau dysregulation in humans^47–49^. Efficient gene knockdown was tested using a pool of short hairpin RNA and its respective scramble control. Our system enabled us to significantly reduce NSun2 protein levels (31.88% decrease, *P* = 0.0018) in the iPSC-derived neuronal cultures (**Supplementary Figure 6**), resembling a similar reduction observed in human cortex of AD brains. Strikingly, using a battery of anti-phospho-tau antibodies (**Figure 2C**), Western blot analysis showed a significant increase in the levels of intracellular phosphorylated tau in several of the phospho-epitopes tested (pSer-199-202, pSer-214, pSer-262, pSer-396-404) upon NSun2 knockdown (**Figure 2D**). In addition, we performed immunostainings on the human iPSC-derived neuronal cultures using the anti phospho-serine 214-tau antibody on the neuronal cultures showing and increase in the number of positive pSer214tau neuronal cells upon NSun2 protein knockdown compared to controls (**Supplementary Figure 7**).

### NSun2 is downregulated in response to AβO

One of the main factors that distinguish AD from other primary tauopathies is the accumulation of AβO species in the brain, which correlates with cognitive decline and/or disease progression in AD patients and animal models^50^. Considering the observed significant reduction of NSun2 levels in the AD brains (**Figure 1**), we next asked whether Aβ could be triggering this pathological alteration. In order to determine if oligomeric Aβ, in a sub-apoptotic concentration (**Supplementary Figure 8**), could promote downregulation of NSun2 levels we exposed rat primary hippocampal neurons from wild type rats to 300nM AβO. This resulted in a progressive decrease of NSun2 protein levels over 24 hours of exposure (**Figure 3A**). Conversely, NSun2 mRNA levels were not affected as a result of the AβO exposure (**Figure 3B**). Concomitantly, at 24 hours we observed a significant increase in phospho-tau levels assessed by western blot using an anti phospho-serine 214 tau antibody (198% increased, *P* = 0.0102; **Figure 3C**). When we performed immunostaining of the neuronal cell culture, consistent with the western blot results, we observed a reduction in the nuclear and dendritic (MAP2 positive) NSun2 signal (**Supplementary Figure 9**). Similarly, to neuronal cultures exposed to AβO, shRNA mediated knockdown of NSun2 protein (**Supplementary Figure 10**) resulted in a significant increase of in the levels of phospho-tau, when tested using antibodies against tau phospho-serine 214 (57.89 % increased, *P* = 0.0216) and tau phospho-threonine 231 respectively (33.82 % increased, *P* = 0.0215) (**Figure 3D, E**). Remarkably, when we performed coimmunostaining on human brain sections of AD patients, we found higher levels of phospho-tau (AT8 immunostaining) in neurons with low NSun2 immunostaining (**Figure 3F**). Taken together, these results suggest that downregulation of NSun2 might explain changes on tau proteostasis observed in AD brains by altering a regulatory post-transcriptional mechanism.

**Figure 3.**
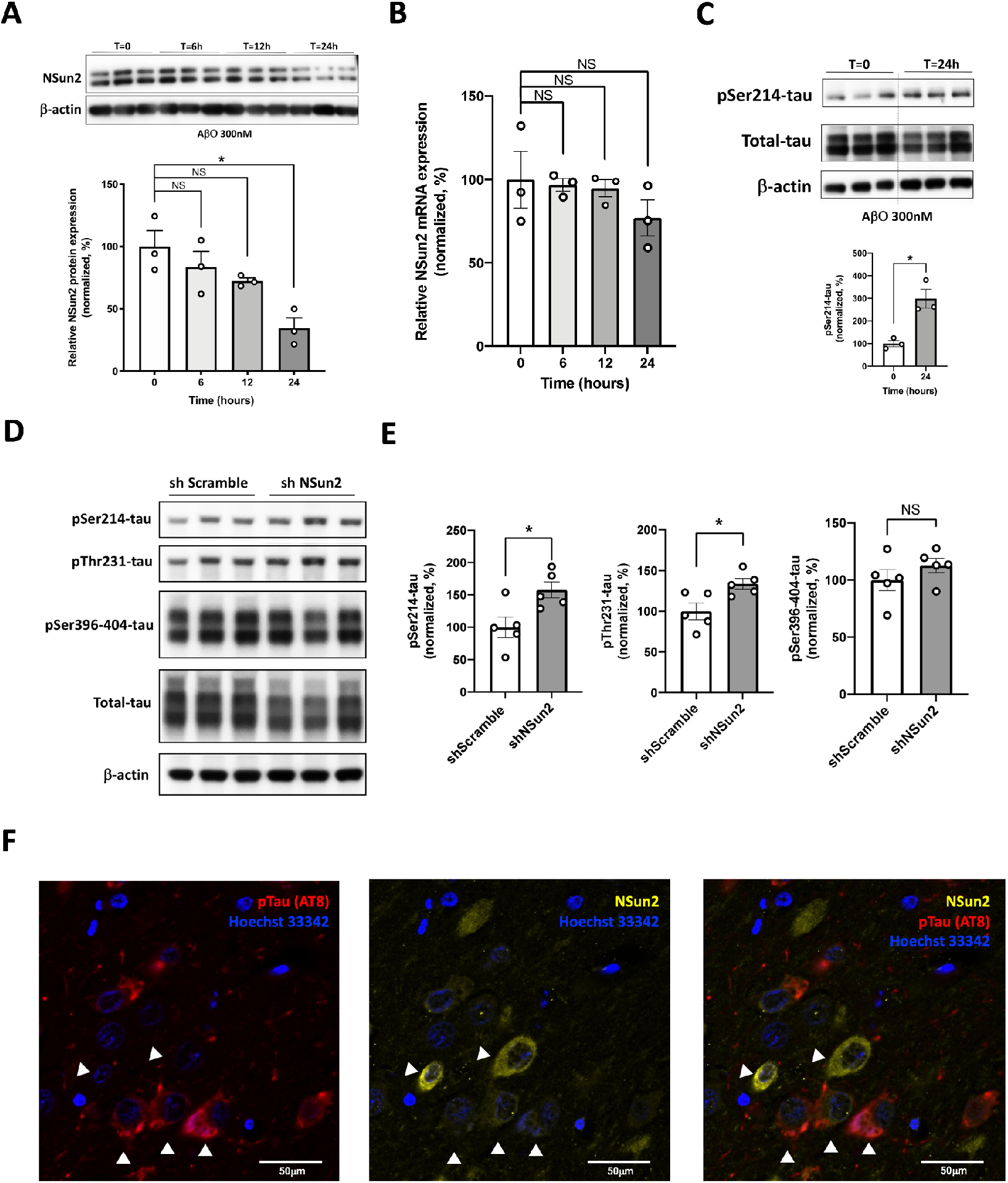
NSun2 is downregulated in response to AβO. (**A**) Western blot quantification of NSun2 protein levels in rat primary hippocampal neurons untreated (control) or treated with 300 nM Aβ oligomers (AβO) for the indicated times (Student’s t test; \**P* < 0.05). Top portion of the panel shows representative Western blots. Histograms show densitometric quantification of NSun2 protein levels (bottom of the panel). NSun2 values are normalized against beta-actin. Data represent mean ± SEM. (**B**) qPCR analysis of NSun2 mRNA levels in rat primary hippocampal neurons untreated (control) or treated with 300 nM AβO for the indicated times. No significant changes in the levels of NSun2 mRNA are found. (**C**) Western blot quantification of phospho-tau levels in rat primary hippocampal neurons untreated (control) or treated with 300 nM AβO for 24 hours. Top portion of the panel shows representative Western blots. Phospho-Serine 214-tau corresponds to the top band; Total tau corresponds to the middle band and beta-actin to the bottom band. Densitometric quantification of phospho-tau protein levels is shown at the bottom of the panel. Phospho-tau levels are plotted relative to the levels of total tau (after normalization of total tau with β-actin levels) (Student’s t test; \**P* < 0.05). (**D**) Rat primary hippocampal neurons were transduced with shNSun2 or scramble control, and protein lysates were collected and analyzed. Representative Western blots with indicated antibodies demonstrated the effects of shNSun2 on the levels of phosphorylated forms of tau. (**E**) Quantification of phosphorylated tau in neurons transduced with shNSun2, plotted relative to the levels of total tau (after normalization of total tau with β-actin levels). (Student’s t test; \**P* < 0.05). Data represent mean ± SEM. (**F**) Double immunofluorescence with NSun2 (yellow) and phospho-tau (AT8, red) antibodies in hippocampus of human AD brain. White arrowheads show NSun2 and AT8 positive immunostaining. Scale bars, 50 μm.

### NSun2 deficiency promotes epitranscriptomic alterations in miR-125b

It has been shown that NSun2 mediates *N*^6^-adenosine methylation (m6A) of miR-125b-5p (abbreviated hereafter as miR-125b), repressing its processing and function^23^. Importantly, miR-125b is found to be upregulated in AD^51–54^ and its upregulation promotes tau hyperphosphorylation and cognitive deficits *in vivo*^55^. To study whether NSun2 deficiency alters m6A methylation of miR-125b and promotes tau hyperphosphorylation *in vivo*, we performed RNA immunoprecipitation and histological analysis. RNA isolated from NSun2 knockout mice brain cortical samples was immunoprecipitated using an anti-m6A antibody and the presence of methylated miR-125b in the immunoprecipitated materials was analyzed by QPCR. In **Figure 4A**, we show the m6A antibody could effectively immunoprecipitate miR-125b. As a negative control, using IgG failed to immunoprecipitate miR-125b. The levels of methylated miR-125b decreased significantly in brain samples from NSun2 deficient mice (43.90 %, *P* = 0.0182) (**Figure 4A**). As expected, miR-125b levels were significantly increased in the input samples from NSun2 knockout mice brains (**Figure 4B**). Similarly, the levels of miR-125b are increase in AD frontal cortex samples (**Supplementary Figure 11**). To further confirm NSun2 deficiency promotes tau alterations *in vivo*, we performed AT8 (phospho-ser-202/Thr205-tau) immunostaining on the brains of aged NSun2 conditional knockout animals. Strikingly, we observed increased immunoreactivity of AT8 positive neurons in the frontal cortex and dentate gyrus of NSun2 deficient mice (right panels, **Figure 4C**). These results support that NSun2 deficiency promotes alterations in m6A methylation of miR-125b and tau hyperphosphorylation.

**Figure 4.**
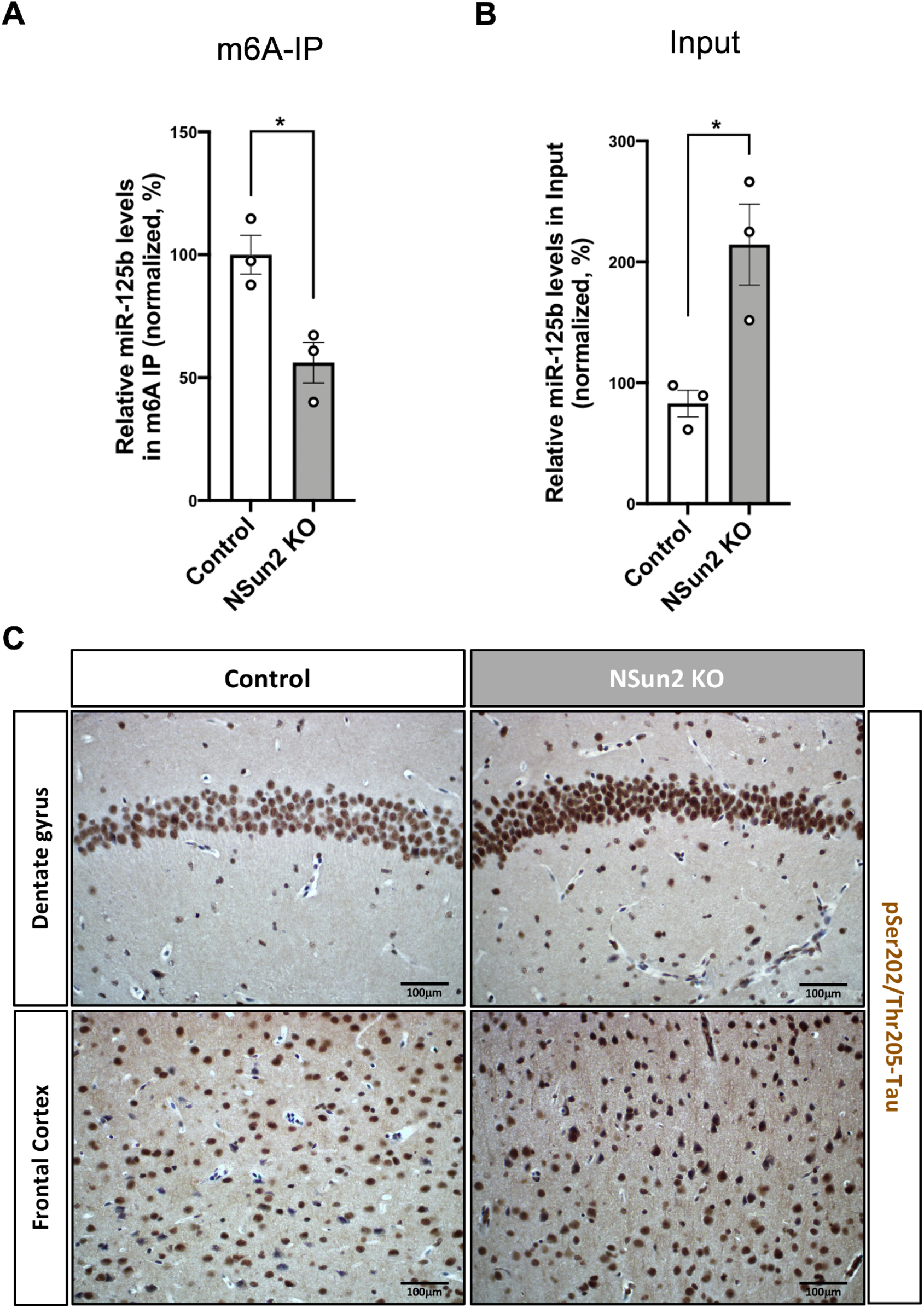
NSun2 deficiency alters miR-125b methylation, miR-125b levels, and promotes tau hyperphosphorylation *in vivo*. (**A-B**) RNA was isolated from the forebrain of NSun2 KO and non-transgenic controls and subjected to IP analysis using anti-m6A or IgG antibody. The presence of miR-125b in the Input (*n*=3) m6A (*n*=3) and IgG (*n* = 3; used as negative control) materials was analyzed by QPCR. Histograms show quantification of miR-125b levels with respect to control in Input and m6A materials. (Student’s *t* test; *\*P* < 0.05). Data represent mean ± SEM. (**C**) Representative images of immunohistochemistry with AT8 antibody in the dentate gyrus (top panels) and frontal cortex (bottom panels) of 9-month-old NSun2 KO and non-transgenic control mice showing a marked increase in phospho-tau immunostaining in NSun2 KO animals. Scale bars, 100 μm.

## Discussion

Our results confirm that NSun2, one of the best described RNA methyltransferases in higher eukaryotes^56–62^, is expressed in the adult human brains (**Figure 1**). However, current understanding of the role that epitranscriptomic regulation plays in brain function and dysfunction is limited^63–65^. NSun2 deficiency has a negative impact on learning and memory in fruit flies and causes intellectual disability and neurological abnormalities in humans^62, 66^. Here, we show NSun2 immunostaining localized in dendrites of neurons in the hippocampal formation and the frontal cortex (**Figure 1**), suggesting epitranscriptomic regulation of neuronal synaptic function is taking place in the adult human brain^67^. Further experimental evidence would be necessary to rule out the role NSun2 plays in synaptic function and regulation. Nevertheless, even though we only observed positive NSun2 staining in neurons, it should be recalled that NSun2 mRNA is also found in the transcriptomic profile of other cell types in the brain^68, 69^, reinforcing NSun2’s role during development and in cellular stress responses^19^.

To date, several studies have implicated epitranscriptomic regulation of coding and non-coding RNA in diverse biological functions^65, 70^. Here, we show NSun2 RNA methyltransferase protein levels are decreased in AD brains, denoting epitranscriptomic alterations in the AD process (**Figure 1**). Unexpectedly, our results show that NSun2 protein is not reduced in PART and PSP cases indicating a distinct disease mechanism in the AD brains. Future studies will be important to uncover whether NSun2 alterations are occurring in specific phases or throughout AD progression. Similarly, further characterization of epitranscriptomic changes and the regulatory proteins of the epitranscriptome (including writers, erasers and readers) in AD, related tauopathies, and other neurodegenerative disorders is warranted.

It has been shown NSun2 methylates miR-125b repressing its processing and function^23^. Here, we have confirmed miR-125b is upregulated in AD brains in the prefrontal cortex and our results in a *Drosophila* model of tau toxicity supports a neuroprotective role for NSun2. In addition, we have shown NSun2 deficiency promotes alterations in methylated miR-125b levels and modulates tau proteostasis *in vivo* using the a NSun2 conditional knockout mouse model. However, it has been described that the loss of NSun2 results in the accumulation of tRNA fragments^71^. Curiously, tRNA fragments function as short RNAs with multi-faceted roles in disease processes^72, 73^, including neurological disorders and possibly AD^74–78^. Therefore, further analysis will be required to uncover other salient roles of NSun2 on the post-transcriptional regulation of non-coding small RNAs in AD and related neurological disorders.

Given that our data supports that NSun2 reduction occurs in response to Aβ accumulation and leads to tau proteostasis alterations, our study finds preliminary evidence that NSun2 targeting could be of therapeutic value. One plausible way of modulating NSun2 could be through Proteinase activated-receptor 2 (PAR2), one of the Proteinase-activated receptors with profound roles in the nervous system^79^. PAR2 has been shown to modulate NSun2 and miR-125b methylation^80^. Moreover, PAR2 expression is reduced *in vivo* in response to Aβ. Furthermore, PAR2 receptor levels are found reduced in human AD brains^81^ which could potentially explain our observed alterations in NSun2.

In conclusion, our results suggest that tau toxicity is modulated by Nsun2 through regulation of tau phosphorylation. This conclusion is based on the fact that NSun2 deficiency regulates tau phosphorylation *in vitro* and *in vivo* and tau toxicity is bidirectionally regulated by NSun2 overexpression and inhibition *in vivo*. Our findings are consistent with NSun2 influencing tau phosphorylation at the post-transcriptional level, perhaps through microRNA regulation, but this may differ depending on the experimental context. Future studies to validate role of NSun2 in other disease models and pertinent behavioral studies would be valuable. It is unlikely that NSun2 is the only methyltransferase with the ability to regulate microRNAs, but the role of NSun2 is of critical interest, given the extraordinary and unique roles this enzyme plays in physiology and pathology. Further investigation will provide a better understanding of tau regulation and advance our understanding of the pathogenesis of neurofibrillary degeneration which might bring us a step closer to the development of novel therapeutic strategies.

### Study approval

The studies using de-identified *post-mortem* autopsy tissue were reviewed and approved by the Columbia University institutional review board (IRB) (New York, NY). *Drosophila* studies are not subject to IRB oversight.

## Author contributions

I.S.M and Y.K conceptualized the project and designed the study methodology. Y.K., J.B., T.W., A.K., E. T and A.A.S., performed research; A.A.S., A.F.T, F.B., S.A. and G.H. contributed resources (study materials, iPSC derived neuronal cultures, *post-mortem* brain samples). A.A.S., A.F.T, F.B., S.A., G.H., and I.S.M supervised or managed the research. Y.K. and I.S.M. analyzed, interpreted and visualized the data; Y.K. and I.S.M. wrote the initial manuscript paper; All authors provided critical feedback and contributed to the final manuscript.

## Acknowledgments

This work was supported by NIH grants R01NS095922 and P50AG0008702 to I.S.M., R01MH117790 to S.A., R03NS112785, R21AG070414-01 and K08NS116166-01 to G.H., and R01AG050658 to F.B. and NIMH Postdoctoral fellowship F32MH115565-01A1 to J.B. Additional support was provided to I.S.M. by the Alzheimer’s Association (NIRG-15-3644-58). A.A.S. is supported by the Henry and Marilyn Taub Foundation and the Thompson Family Foundation Program (TAME-AD). We want to thank Jean Paul Vonsattel for neuropathology support. We are grateful to the late Dr. Peter Davies for his generous gift providing us PHF1 anti-phospho tau antibody. Finally, we want to thank Prof. Dr. Stephan J. Sigrist who kindly shared with us the transgenic *Drosophila* line.

