## Supplementary materials for "NSun2 deficiency promotes tau hyperphosphorylation and neurodegeneration through epitranscriptomic regulation of miR-125b"

**This PDF file includes**

Supplementary Methods

Supplementary Figures 1 to 11

Supplementary Tables 1 to 2

Supplementary  References

**Methods**

*Human brain tissue samples*. Autopsy brain samples were obtained from the New York Brain Bank at Columbia University Medical Center. The demographics of human cases identified in the Columbia University Alzheimer's Disease Research Center Neuropathology Core and used in this study are listed in detailed in **Supplementary Table 1**. These specimens were obtained by consent at autopsy and have been de-identified and are IRB exempt so as to protect the identity of each patient. Formalin-fixed paraffin embedded (FFPE) specimens were sectioned by the Histology Service at Columbia University Medical Center. Immunohistochemistry was performed on 6 μm paraffin-embedded sections as previously described (*1*) using various antisera (**Supplementary Table 2**). Images were captured using an Olympus BX53 microscope with an Olympus camera DP-72 (Olympus Lifescience).

*Mice*. All animal studies were performed according to protocols examined and approved by the Animal Use and Care Committee of Columbia University and Icahn School of Medicine at Mount Sinai. The J20 transgenic mouse line expressing a mutated human APP (hAPP: K670N/M671L and V717F) under the control of the platelet-derived growth factor promoter was obtained from The Jackson Laboratories (Stock # 034836). C57BL/6N-Nsun2$^{tm1c(EUCOMM)Wtsi}$/WtsiOulu mutant mouse sperm were obtained from the Wellcome Trust Sanger Institute (Cambridgeshire, UK). Mice with the tm1c allele exhibit a phenotypically wild-type state although exon 6 of Nsun2 is flanked by loxP sites, where presence of Cre-recombinase excises this exon and produces a frame-shift, resulting in early termination of NSun2 translation. *In vitro* fertilization was performed with C57BL/6N-Nsun2$^{tm1c(EUCOMM)Wtsi}$/WtsiOulu mutant mouse sperm and wildtype (control) C57BL/6N females. Mice were then bred further to obtain *Nsun2*$^{2lox/2lox}$ mice (2 copies of tm1c mutant allele). *Nsun2*$^{2lox/2lox}$ mice were crossed with a *CamK-Cre*$^{+}$ line (*2-5*) to produce *CamK-Cre*$^{+}$,*Nsun2*$^{2lox/2lox}$ mice for knockout of *Nsun2* in excitatory forebrain neurons. Immunohistochemistry was performed on 7 μm paraffin-embedded sections as previously described (*6*) using various antisera (**Supplementary Table 2**). Images were captured using an Olympus BX53 microscope with an Olympus camera DP-72 (Olympus Lifescience).

*Drosophila stocks*. All *Drosophila* stocks were maintained on standard food (Bloomington recipe, Archon Scientific) in incubators at constant 70% relative humidity and 25°C on a 12-h/12-h light/dark cycle. *Drosophila* dNSun2 line was generated by Prof. Dr. Stephan J. Sigrist (*7*), who kindly shared with us. EGFP control line (Stock #5430), NSun2-RNAi line (Stock #62495) and GFP-RNAi control line (Stock #42555) were obtained from Bloomington *Drosophila* Stock Center. For tau overexpression, we used the *Drosophila* line previously generated by us (*8*) with the full-length human tau coding sequence (2N4R). The GMR-GAL4 driver was used for expression of transgenes in the eye.

*Drosophila rough eye phenotype assessment.* For light microscopy imaging of the *Drosophila* eyes, 7-day old adult flies were collected and eye images recorded, in a blinded fashion, by three independent observers as described by others with slight modifications (*8, 9*). Briefly, fly eye images of specified genotypes were acquired under a dissecting microscope by one researcher, coded and given to another two researchers for blind quantitative assessment. To accurately quantitate eye degeneration, we used an Olympus Stereoscope SZX16 microscope in combination with the QCapture Pro7 Imaging software. The extent of eye area undergoing degeneration was analyzed using the image processing and analysis program Image J (Image J, NIH) following previously described methods (*10*, *11*).

*Cell culture*. HEK 293T cell line was used for optimal lentivirus production. HEK 293T cells were obtained from the American Type Culture Collection were grown either in Dulbecco's modified Eagle's medium (Thermo Fisher Scientific) supplemented with 10% fetal bovine serum, 2 mM glutamine, 100 units/ml penicillin, and 100 μg/ml streptomycin in a humidified atmosphere of 5% CO2 and 95% air at 37 °C. Human induced pluripotent stem cells (hiPSCs) knockin with the APP London mutation (V717L) were generated by Dr. Andrew Sproul's laboratory using CRISPR/Cas9 on the control IMR90 cl.4 iPSC line (WiCell) (*12*). Human $APP^{V717L}$ iPSC and isogenic controls were maintained feeder-free in StemFlex media (Thermo Fisher Scientific) and Cultrex substrate (Bio-techne). Transdifferentiation of human $APP^{V717L}$

iPSC and isogenic controls into cortical-like pyramidal neurons was performed following established protocols (*13*). Alternatively, neural progenitor cells were generated from iPSC controls, cultured and differentiated into neurons following established protocols (*14*).

Primary hippocampal neuronal cultures were prepared following previously established methods with slight modifications (*15, 16*). Briefly, hippocampi were dissected from E18 rats, and neurons plated on 100 µg/mL poly-D-lysine-coated 12-well-plates at the density of $1.5x10^5$ cells/well for biochemistry assays, or $6x10^4$ cells/coverslip on 18 mm coverslips for immunofluorescence. Primary neurons were maintained in Neurobasal medium (Thermo Fisher Scientific) with the B-27 supplement (Thermo Fisher Scientific) and 0.5 mM glutamine (Thermo Fisher Scientific) at 37°C, and 1/3 of the medium was changed every 3-4 days up to 3 weeks in culture.

*Lentiviral shRNA preparation*. The human pull of shRNA NSun2 and scramble shRNA control were obtained from Applied Biological Materials Inc. For lentivirus preparation for *in vitro* experiments, the second-generation packaging system (which generates replication-deficient lentivirus) was used for all experiments (*17*). Packaging vectors psPAX2 and pMD2.G were obtained from Addgene. Briefly, lentiviral constructs for shRNA or scramble control were co-transfected with the packaging vectors into HEK293T cells using CalFectin (Signagen). Supernatants containing virus were collected 48 h after transfection. After centrifugation at 1000 rpm for 10 min, the supernatants were passed through a 0.45 µm PDVF filter unit (Nalgene). The viruses were concentrated 20–30x by centrifugation in an Amicon Ultra centrifugal filter (100 K) (Millipore) following the manufacturer's instructions. Lentiviral stock titration was carried out using the Global UltraRapid Lentiviral Titer Kit (System Biosciences). Viruses were aliquoted and stored at −80°C.

*Amyloid β oligomers preparation*. Oligomer-enriched preparations of Aβ (AβO) were obtained according to previously published methods (*18*). Briefly, lyophilized Aβ42 peptide (rPeptide) was equilibrated to room temperature for 30 min to avoid condensation upon opening the vial. The lyophilized peptide was resuspended in hexafluoro-2-propanol (HFIP; Sigma-Aldrich) to a concentration of 1 mM to allow monomerization for another 2 h at room temperature and then aliquoted into low protein–binding Eppendorf tubes. HFIP was removed by speed vacuum, and the monomers were stored in −80°C. To prepare oligomer-enriched preparations, the aliquots were resuspended in anhydrous DMSO to make a 5 mM solution followed by 10 min of sonication. The resuspended peptide was diluted to 100 µM in ice-cold Ham's F-12 medium, immediately vortexed for 30 s, and then incubated at 4°C for 24 h before use. Total Aβ concentration was measured by bicinchoninic acid protein assay (BCA) after oligomerization, and indicated concentration of oligomeric Aβ was used (see **Figure 3**). For control experiments, vehicle treatment corresponded to the same volume of DMSO and F12 media used for AβO treatment was processed as for Aβ42 oligomerization.

*Tissue and cell lysates preparation for Western blotting*. Samples from human brain were stored at −80 °C and ground with a mortar in a frozen environment with liquid nitrogen to prevent thawing of the samples, resulting in tissue powder. Mouse brains were quickly dissected on an ice-cold plate and the different structures stored at −80 °C. Human and mouse protein extracts were prepared by homogenizing brain structures in ice-cold extraction buffer [250 mM sucrose, 20 mM Tris–HCl (pH 7.4), 1 mM ethylenediaminetetraacetic acid (EDTA), 1 mM ethylene glycol tetraacetic acid (EGTA)] containing a cocktail of protease and phosphatase inhibitors (Halt Protease & Phosphatase Inhibitor Single-Use Cocktail ; Thermo Scientific) using 20 strokes with a Teflon-coated pestle. Homogenates were centrifuged at 3,000 rpm for 5 min at 4 °C. The resulting supernatant was collected, and protein content determined by BCA (Pierce, Rockford, IL, USA).

To prepare cell lysates for Western blotting neurons were collected in 1X Laemmli sample buffer (150ml/well on 24 well plates; Bio-Rad Laboratories). Cells were lysed in and boiled for 5 minutes. A volume of 15 ml was subsequently used for electrophoresis.

*Western blotting*. 10 μg of total protein was resolved by sodium dodecyl sulphate-polyacrylamide gel electrophoresis (SDS-PAGE) and transblotted using standard procedures. Nitrocellulose membranes (BioRad) were blocked in TBS-T (150 mM NaCl, 20 mM Tris–HCl, pH 7.5, 0.1% Tween 20) supplemented with 5% non-fat dry milk. Membranes were incubated overnight at 4 °C with designated primary antibodies (see Table S2) in TBS-T supplemented with 5% non-fat dry milk, washed with TBS and next incubated with HRP-conjugated secondary antibodies (Kindle Biosciences; Cell Signaling Technology) for 1 hour. Afterwards, the membrane was washed with TBS-T and developed using the chemiluminescence ECL kit (Millipore Classico or Kindle Biosciences ECL kit) and imaged using a Kwik Quant Imager (Kindle Biosciences).

For western blotting of Aβ oligomers, 10-20% Tris-Tricine SDS-PAGE was performed. Synthetic Aβ oligomers were prepared in 1X Laemmli sample buffer (Bio-Rad Laboratories) without reducing agent and resolved by SDS-PAGE. The separated proteins were transferred onto Immobilon-FL PVDF membrane (EMD Millipore), and subsequently blocked and incubated with 6E10 monoclonal antibody and secondary antibodies (see **Supplementary Table 2**) as indicated above for nitrocellulose membranes.

*Cell viability assay*. Rat primary hippocampal cultures were treated with AβO preparations and cell viability determined using the CellTiter-Glo Luminescent Cell Viability Assay (Promega) following the manufacturer's instructions.

*RNA isolation*. Fresh-frozen pulverized brain tissue or rat primary hippocampal neurons were lysed in QIAzol and homogenized using a QIAshredder column. Total RNA was extracted using the miRNeasy Kit (Qiagen). RNA concentration and purity were assessed by measuring the optical density at 260 and 280 nm with a Nanodrop Spectrophotometer (Thermo Scientific).

*RNA immunoprecipitation.* For immunoprecipitation of RNA, two rounds using 5 μg of anti-m6A antibody and 4 μg of small RNA were performed. The reaction was carried out using the Immunoprecipitation Kit—

Dynabeads Protein G (ThermoFisher) following previously established protocols with slight modifications (*19*). First, the anti-m6A antibody (**Supplementary Table 2**) was coupled to Dynabeads Protein G in 500 μl of Binding and Washing Solution for 3 hours at 4°C followed by 10 minutes incubation at room temperature. Beads were then washed three times in Washing Buffer. Small RNA was added to the antibody-coupled beads in 1X IP buffer (10 mM Tris-HCl, 150 mM NaCl and 0.1% (vol/vol) Igepal CA-630) supplemented with RNase inhibitor—RNAsin Plus (Promega) and kept on the rotating platform at 4°C overnight. On the next morning beads were washed 4 times with IP buffer. Finally, the beads were resuspended in 100 μl of $m^6A$ competitive elution buffer (1X IP buffer containing 6.7mM $N^6$-Methyladenosine, 5′-monophosphate sodium salt (Sigma-Aldrich, cat. no. M2780) and RNAsin Plus) with continuous shaking for 1 hour at 4 °C. The mixture was placed on a magnetic separation rack, and supernatant containing the eluted $m^6A$ RNA was collected to a new tube. Then, another 100 μl of $m^6A$ competitive elution buffer was added for one more elution. Immunoprecipitated RNA was recovered using the miRNeasy Kit (Qiagen) following the instructions from the manufacturer. As a control, immunoprecipitation was performed using IgG instead of anti-m6A antibody. The rest of experimental parameters were kept identical.

*Quantitative real-time PCR*. *c*DNA synthesis was performed using the First Strand cDNA Synthesis Kit (Origene) following the manufacturer's instructions. Quantitative real-time PCR (QPCR) was performed using *Power* SYBR Green PCR Master Mix (Applied Biosystems) and an Eppendorf Realplex Mastercyler with the following settings: 1 cycle at 95 °C for 10 min and 45 cycles of amplification: 95°C for 30s, 57°C for 1 min and 60°C for 30sec. The following primers pairs were used: human Nsun2 Fwd 5'-ATGATCGAATTTTATGTGATGTCC-3' and human Nsun2 Rvrs 5'-AGGTGGTCCACTTTTTCCAA-3'; human GAPDH 5'-CTGCACCACCAACTGCTTAG-3' and human GAPDH Rvrs 5'-GTCTTCTGGGTGGCAGTGAT-3'; rat Nsun2 Fwd 5'-GTCAGCGAGACGGAGTCTG -3' and rat Nsun2 Rvrs 5'- TGGGGGAATCATGCTGAC -3'; rat GAPDH 5'- GGCAAGTTCAATGGCACAGT -3' and rat

GAPDH Rvrs 5'-TGGTGAAGACGCCAGTAGACTC -3'. The amount of Nsun2 was quantified and normalized to GAPDH mRNA using the comparative CT method.

TaqMan MicroRNA assays were used to measure miR-125b-5p levels (ThermoFisher). 10 ng of total RNA was reverse-transcribed using specific stem-loop reverse transcription primers (ThermoFisher) and miR-125b-5p levels were measured on a Mastercycler ep realplex (Eppendorf). The levels of U6snRNA were used as endogenous controls for normalization using the comparative CT method.

*Immunofluorescence*. For immunofluorescence of human brain sections, human FFPE sections were deparaffinized with Histoclear and rehydrated before performing antigen retrieval. Antigen retrieval procedure was done using Citrate buffer (Biogenex). FFPE sections were blocked in TBST containing 10% normal goat serum for 1 hour at room temperature and incubated with primary antibody (**Supplementary Table 2**) in TBST containing 10% normal donkey serum at 4°C overnight. After three washes with TBST, the tissues were incubated with Alexa Fluor 488 goat anti-rabbit or Alexa Fluor 594 goat anti-mouse IgG (**Supplementary Table 2**) for 3 hours at room temperature. Following three washes with TBS, autofluorescence was quenched with 0.3% Sudan black (Decon Laboratories) in 70% ethanol (Fisher Scientific) for 20 mins at room temperature. The sections were rinsed in TBS. The nuclei were stained Hoechst33342 staining dye solution (ThermoFisher) in TBS for 8 min at room temperature. Following three washes with TBS, sections were mounted on slides using SlowFade gold anti-fade reagent (ThermoFisher) and imaged using confocal laser scanning microscopy via Z stack.

For immunofluorescence of human iPSC derived neurons, we follow previously established methods (*14*). Briefly, iPSC derived neurons on coverslips were fixed with 4% paraformaldehyde (Electron microscopy) and permeabilized with 0.2% in TBS. Coverslips were quenched with ammonium chloride for 10 minutes then blocked with 10% normal goat serum for 30 minutes. Next, coverslips were incubated primary antibodies for NSun2, phosphoSer214 tau and MAP2 (**Supplementary Table 2**). Anti-mouse, anti-rabbit, and anti-chicken secondary antisera conjugated with Alexa Fluor dyes (ThermoFisher-Life Technologies) were used for secondary immunodetection (**Supplementary Table 2**). Finally, coverslips were stained with

DAPI for 10 minutes and mounted using Shandon ImmuMount mounting medium (Fisher Scientific). Labeled neurons were imaged using confocal laser scanning microscopy via Z stack.

For immunofluorescence of rat primary neuronal cultures, cells in coverslips were fixed with 4.0% paraformaldehyde (Electron microscopy) and blocked with 10% normal goat serum in TBS (Thermo Scientific) and incubated overnight in NSun2 antisera (Thermofisher Scientific), AT8 antisera (Thermofisher Scientific) and MAP2 antisera (Abcam). Anti-rabbit, anti-mouse and anti-chicken secondary antisera conjugated with Alexa Fluor dyes (Life Technologies) were used for immunodetection. Labeled neurons were imaged in a Zeiss confocal laser scanning microscope (LSM800).

*Statistical analysis*. For quantitative immunoblotting, rough eye phenotype assessment, and QPCR experiments, the statistical significance was determined by Student's t test or Mann-Whitney *U* test using GraphPad Prism (GraphPad Software, La Jolla, CA, USA). A P value of less than 0.05 was considered significant. For all figures in which error bars are shown, data represent the mean ± SEM. Statistical outliers and specimens with measurement errors were excluded.

*Study approval*. Studies using autopsy tissue were approved by the IRB of Columbia University. Written informed consent was provided by the next of kin. *Drosophila* studies are not subject to IRB oversight.

**Supplementary Figures**

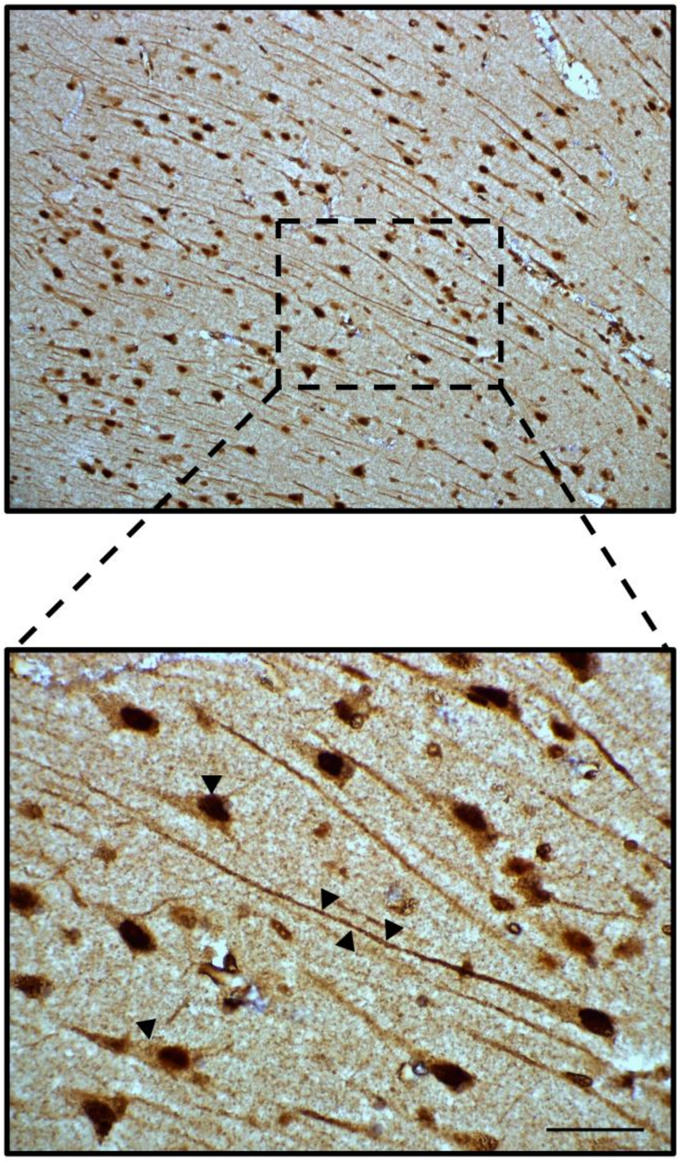

**Supplementary Figure 1. Subcellular localization of neuronal NSun2 protein in the human brain**. Representative immunohistochemistry performed in formalin-fixed paraffin embedded sections of a human control brain using a human specific NSun2 antibody. Black arrowheads show NSun2 distribution in the nucleus, soma, and neurites of neurons in the CA1 region of the hippocampus (magnification in bottom panel). Scale bar, 50 µm.

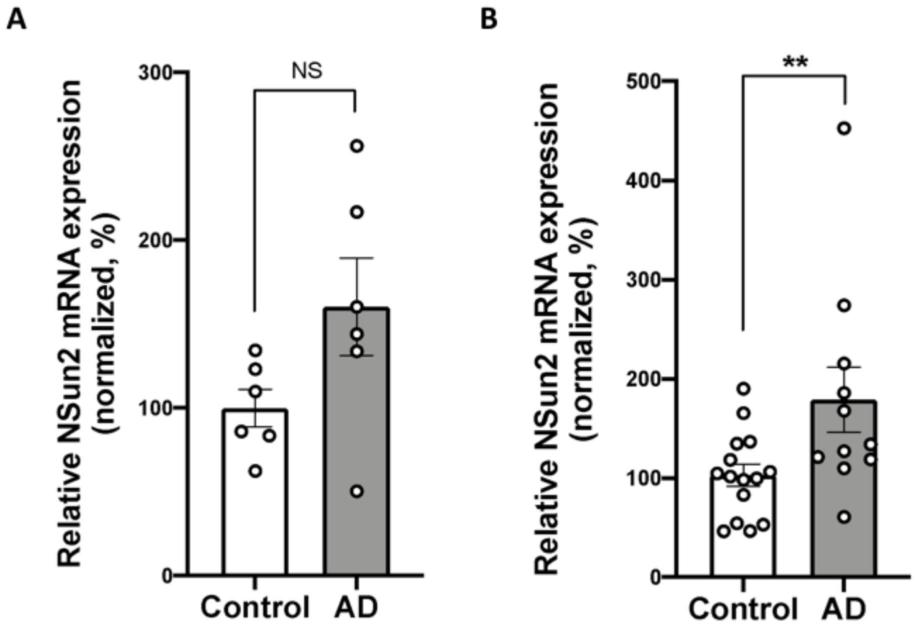


**Supplementary Figure 2. NSun2 mRNA levels in human brains**. (**A**) QPCR analysis shows no difference in the levels of NSun2 mRNA in AD brains ($n$ = 6) compared to control ($n$ = 6) in the hippocampus. (**B**) QPCR analysis of prefrontal cortex samples shows a significant increase of NSun2 mRNA levels in AD ($n$ = 11) brains compared to controls ($n$ = 15) (normalized to GAPDH mRNA) (Mann-Whitney $U$ Test; $^{**}P < 0.01$). Data represent mean ± SEM.

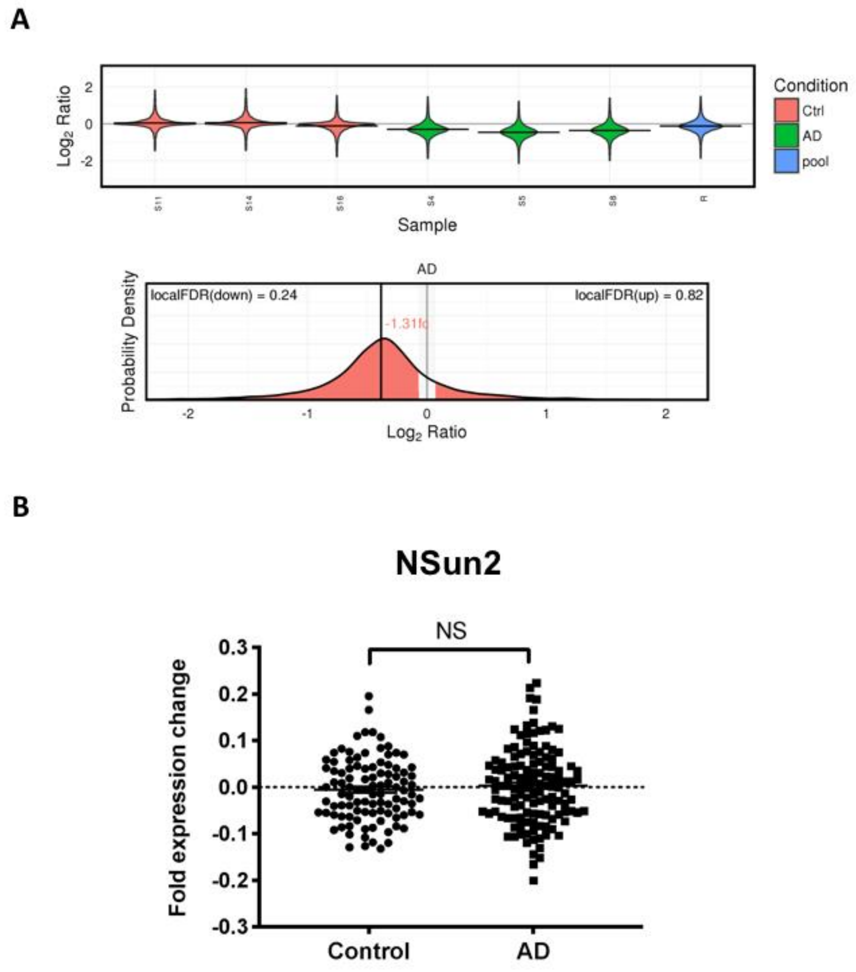


**Supplementary Figure 3. Multigroup comparisons of NSun2 protein and mRNA levels among the AD and control groups**. (**A**) Probability density plots for NSun2 from hippocampus AD proteomic dataset is shown. Algorithm used by Xu et al. (*20*) (http://www.dementia-proteomes-project.manchester.ac.uk/ Proteome/Search) calculates a local FDR (1-p (protein differs from control by at least 5%)) both for upregulation and downregulation. Variability across all hippocampus samples (Alzheimer disease and controls) is shown (top panel). Mean Fold change (fc) is indicated in the AD density plot for NSun2 protein (bottom panel).

(**B**) Histogram representing the NSun2 mRNA fold change expression values from the microarray datasets GSE44772; GSE44768; GSE,44770 and GSE 44771 are shown. No significant change in the levels of NSun2 mRNA was observed. Statistical significance was analyzed by Mann–Whitney *U* test.

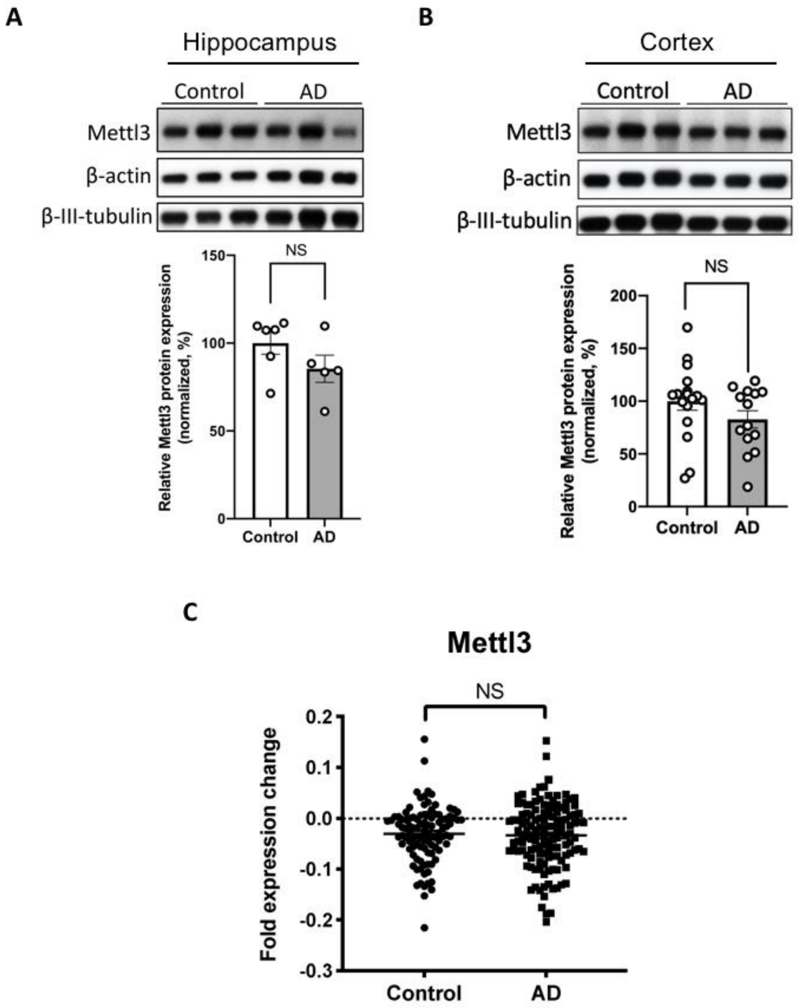


**Supplementary Figure 4. Mettl3 expression in the AD human brain**. (**A**) Western blot quantification of Mettl3 protein levels in human hippocampus of controls (*n* = 6) and AD (*n* = 5). (**B**) Western blot quantification of Mettl3 protein levels in prefrontal cortex of AD patients (*n* = 14) compared to their respective controls (*n* = 17). (**A-B**) Histograms show densitometric quantification of Mettl3 protein abundance with respect to control at the bottom of the panels. Mettl3 is normalized by β-actin in all samples. No significant change in the levels of Mettl3 protein was observed between controls and AD brains. Statistical significance was analyzed by Mann–Whitney *U* test. Data represent mean ± SEM. (**C**) Histogram representing the Mettl3 mRNA fold change expression values from the microarray datasets GSE44772; GSE44768; GSE44770 and GSE44771 are shown. No significant change in the levels of Mettl3 mRNA was observed. Statistical significance was analyzed by Mann–Whitney *U* test.

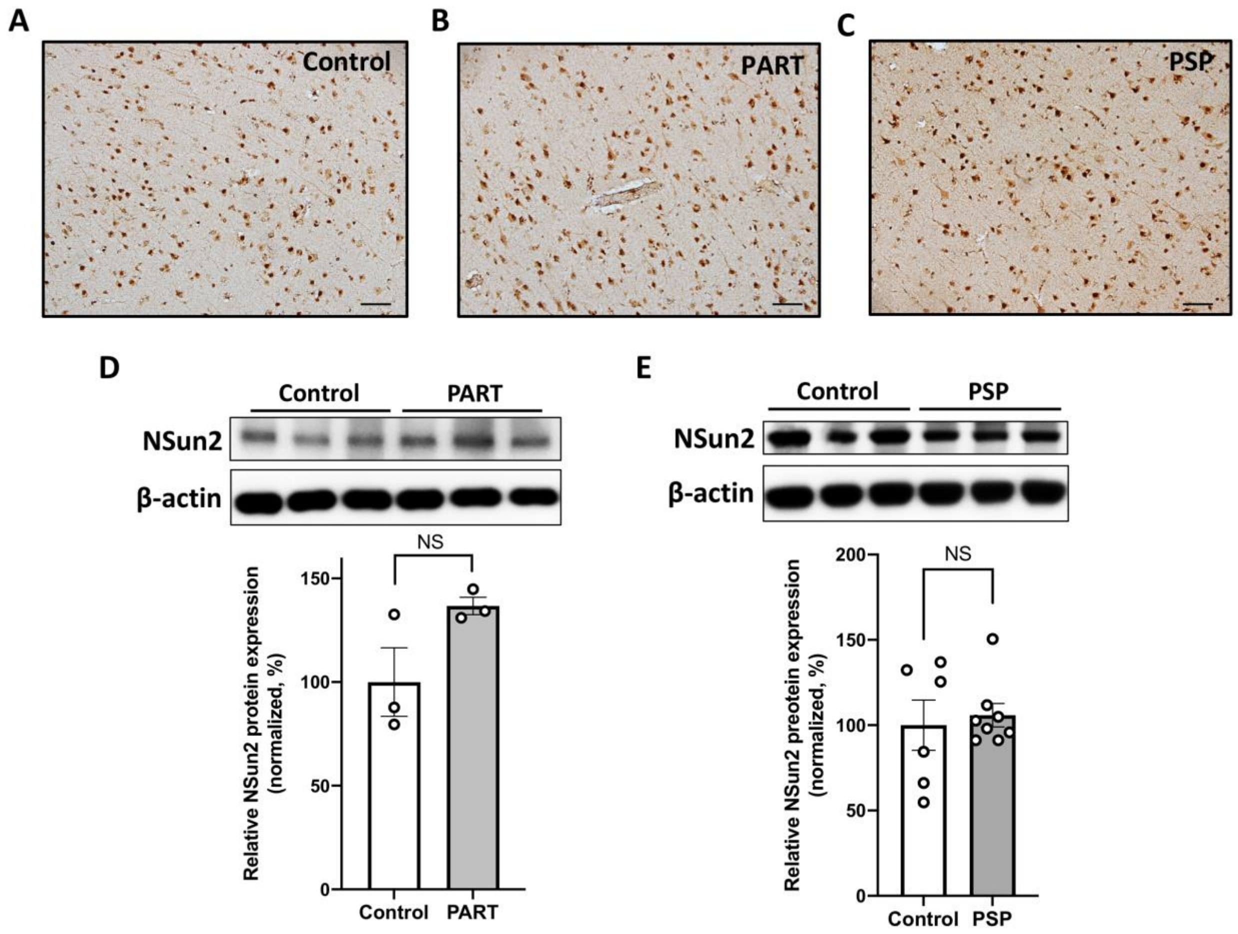


**Supplementary Figure 5. Brain NSun2 protein levels in human tauopathy**. (**A-C**) Shown are representative NSun2 immunohistochemistry images of Primary Age-Related Tauopathy (PART), Progressive Supranuclear Palsy (PSP) and Control human brains. Scale bars, 120 μm. (**D**) Western blot quantification of NSun2 protein levels in human hippocampus of controls (*n* = 3) and PART (*n* = 3). (**E**) Western blot quantification of NSun2 protein levels in Globus Pallidus-putamen of PSP patients (*n* = 8) compared to their respective controls (*n* = 6). (**D-E**) Histograms show densitometric quantification of NSun2 protein abundance with respect to control at the bottom of the panels. NSun2 is normalized by β-actin in all samples. No significant change in the levels of NSun2 protein was observed between controls and PART or PSP brains. Statistical significance was analyzed by Mann–Whitney *U* test. Data represent mean ± SEM**.**

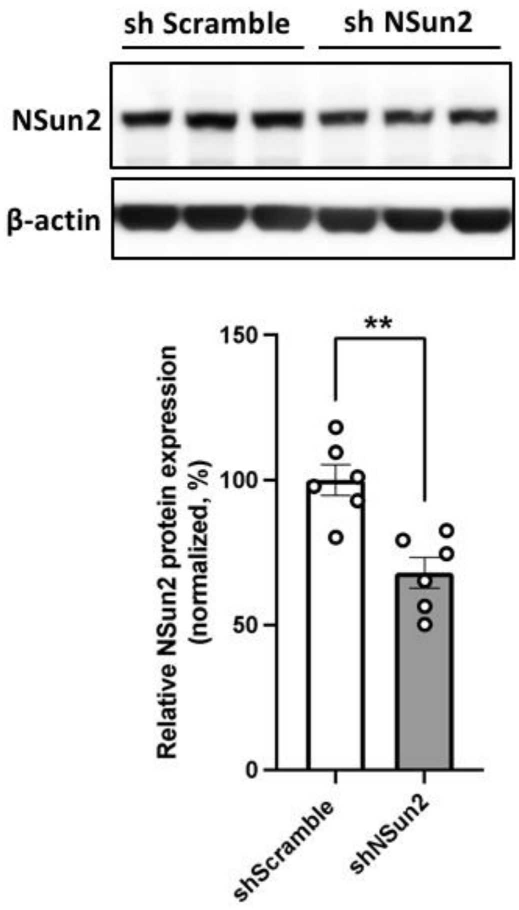


**Supplementary Figure 6. NSun2 knockdown in human iPSC derived neuronal cultures**. Human iPSC derived neurons were transduced with shNSun2 or scramble control, and protein lysates were collected and analyzed. Representative western blots with indicated antibodies demonstrated the effects of shNSun2 on the levels of NSun2 protein (top portion of the panel). Histogram show densitometric quantification of NSun2 protein abundance with respect to control at the bottom of the panel. NSun2 is normalized by β-actin in all samples (Student's t test; $^{**}P < 0.01$). Data represent mean ± SEM.

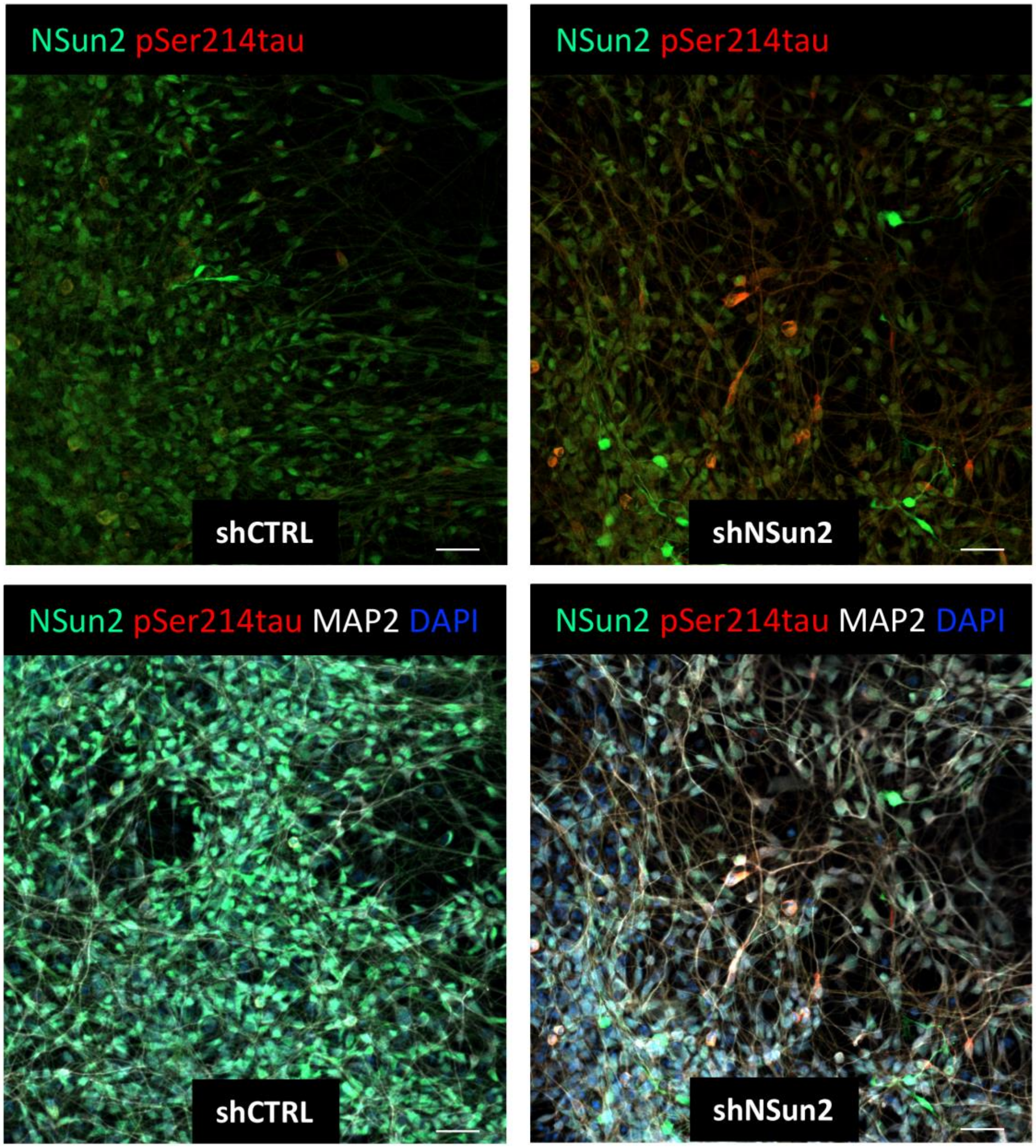


**Supplementary Figure 7. Tau hyperphosphorylation in NSun2 deficient human iPSC derived neurons**. Representative immunofluorescence images of Human iPSC derived neurons transduced with shNSun2 (right panels) or scramble control (shCTRL; left panels) immunostained using NSun2 (green), phospho-serine 214 tau (pSer214tau; red) and MAP2 (white) antibodies. Scale bars, 50 µm.

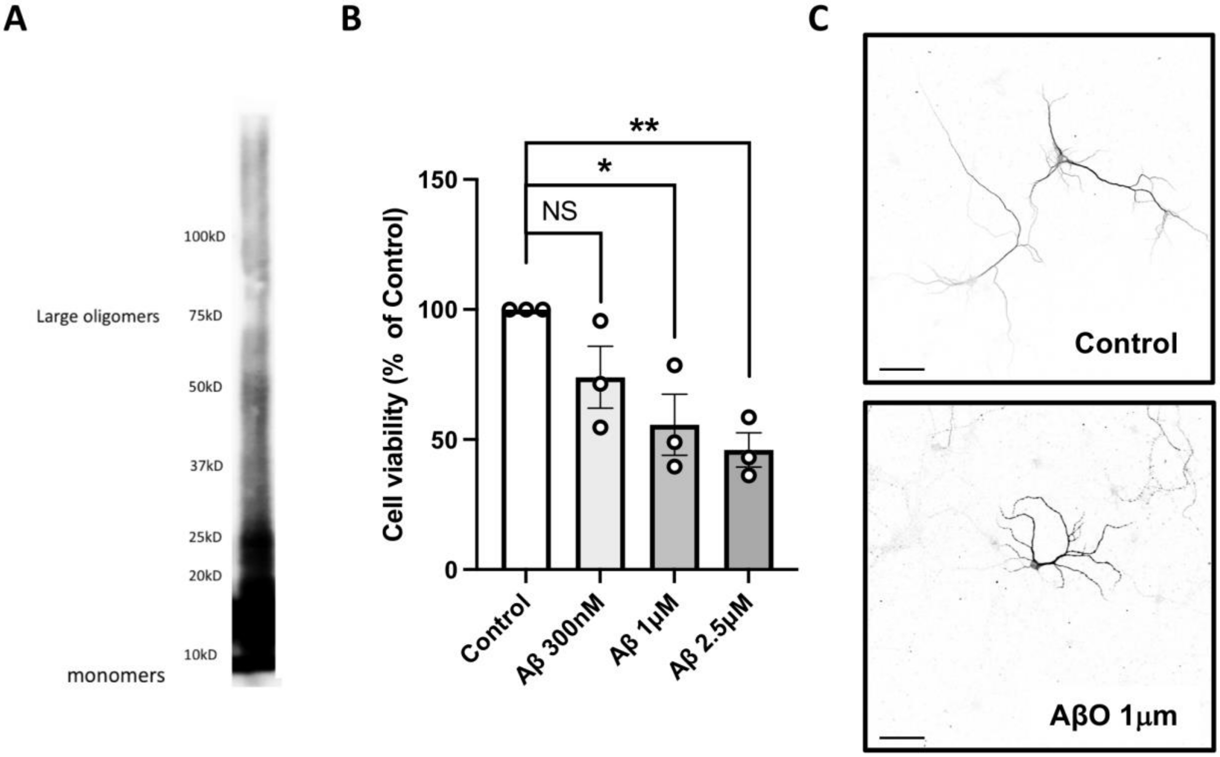


**Supplementary Figure 8. Characterization of amyloid beta preparations**. (**A**) Representative western blot using the anti-amyloid-β (Aβ) antibody 6E10 after incubation for 1 day. Aβ species are categorized as monomers (~4 kDa), oligomers (~8-75 kDa) and fibrils (>75 kDa) based on their molecular weight (*18*). (**B**) Time course of cell death elicited by Amyloid beta oligomers (AβO) treatment. Cell viability assay shows the percentage of remaining viable neuronal cells in cultures vehicle-treated (control) or treated with 300nM–2.5 μM for 24 h. Cell death becomes evident by 24 h with 1 μM and 2.5 μM when compared to controls (Student's t test; $*P < 0.05$; $**P < 0.01$). Data represent mean ± SEM. (**C**) MAP2 immunofluorescence revealing altered dendritic morphology after AβO exposure compared to control treatment. Scale bars, 50 μm.

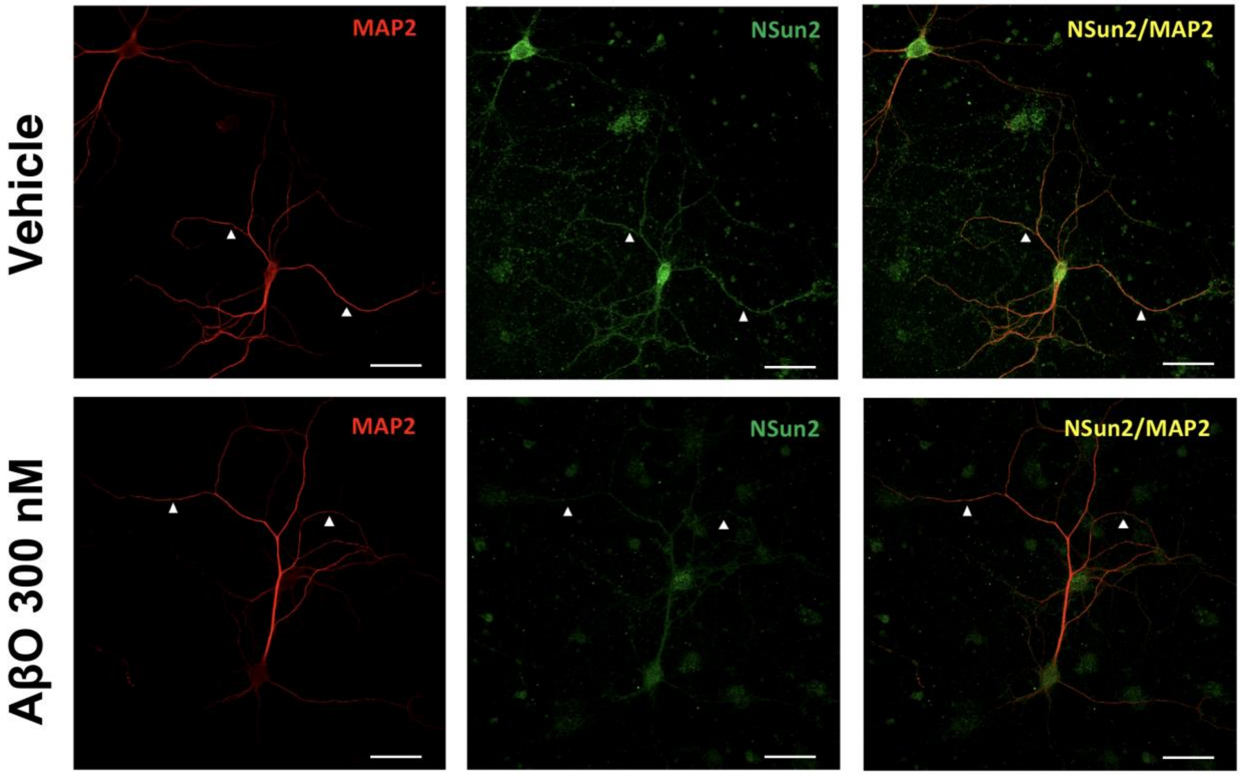


**Supplementary Figure 9. Effect of AβO on NSun2 levels and localization in rat primary neuronal cultures**. Representative immunofluorescence images of vehicle-treated (top panels) and AβO-treated (bottom panels) rat primary neuronal cultures stained for Microtubule associated protein 2 (MAP2; red, dendritic marker) and NOP2/Sun RNA methyltransferase 2 (NSun2; green). Images examination reveal that NSun2 was present in dendrites, similarly observed in human neurons (**Figure 1**) (top panels, indicated with white arrowheads). Colocalization of proteins is represented in yellow. Representative immunofluorescence images of cultured hippocampal primary neurons (Day in vitro 21) treated for 24 h with AβO 300 nM and stained for MAP2 (Red), and NSun2 (green) is shown in the bottom panels. In the presence of 300 nM AβO, NSun2 clearly decreases in dendrites (indicated with white arrowheads) and the nucleus.

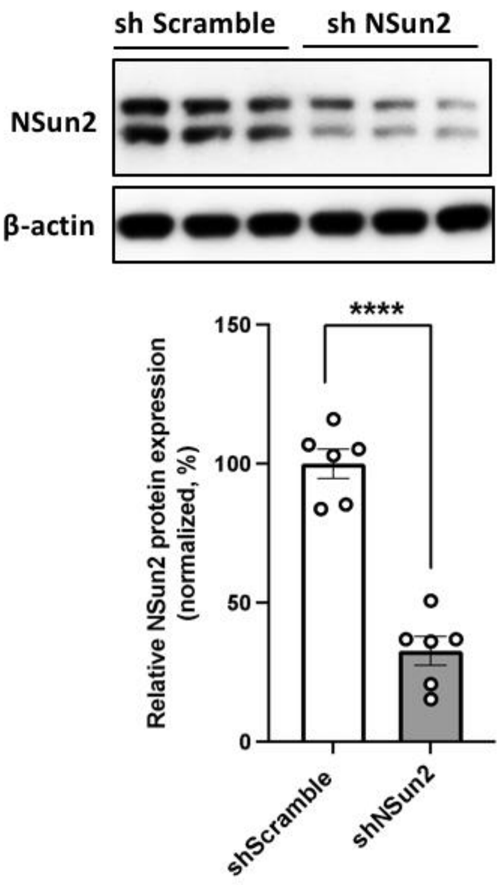


**Supplementary Figure 10. NSun2 knockdown in rat primary neuronal cultures**. Rat primary neurons (Day *in vitro* 21) were transduced with shNSun2 or scramble control, and protein lysates were collected and analyzed. Representative Western blots with indicated antibodies demonstrated the effects of shNSun2 on the levels of NSun2 protein (top portion of the panel). Histogram show densitometric quantification of NSun2 protein abundance with respect to control at the bottom of the panel. NSun2 is normalized by β-actin in all samples (Student's t test; ****$P < 0.0001$). Data represent mean ± SEM.

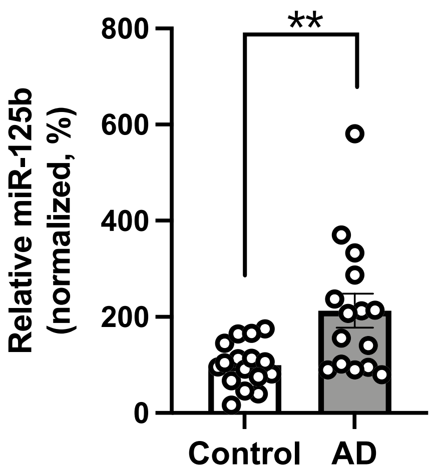


**Supplementary Figure 11. miR-125b levels in human AD brains**. QPCR analysis of prefrontal cortex samples shows a significant increase of miR-125b levels in AD (*n* = 15) brains compared to controls (*n* = 16) (normalized to U6 small nuclear RNA) (Mann–Whitney *U* test; $**P < 0.01$). Data represent mean ± SEM.

### Supplementary Tables

**Supplementary Table 1. Case demographics**

| Arbitrary Case # | Gender | Age at Death (yr) | PMI (hr) | NPDX | Braak stage |
|---|---|---|---|---|---|
| 1 | M | 95 | 3.25 | Control | III-IV |
| 2 | M | 67 | 33.66 | Control | 0 |
| 3 | F | 82 | 3.8 | Control | I-II |
| 4 | M | 80 | NA | Control | 0 |
| 5 | M | 90 | NA | Control | I-II |
| 6 | M | 81 | 5.88 | Control | V-VI |
| 7 | M | 93 | NA | Control | I-II |
| 8 | M | 77 | 3.16 | Control | III-IV |
| 9 | M | 90 | 2.16 | Control | III-IV |
| 10 | F | 90 | 5.41 | Control | III-IV |
| 11 | F | 101 | 3.45 | Control | III-IV |
| 12 | F | 84 | 1.41 | Control | III-IV |
| 13 | F | 84 | NA | Control | I-II |
| 14 | F | 85 | 5.53 | Control | I-II |
| 15 | F | 68 | 7.25 | Control | 0 |
| 16 | F | 92 | 2 | Control | III-IV |
| 17 | M | 92 | NA | Control | I-II |
| 18 | M | 87 | 5.16 | Control | III-IV |
| 19 | M | 94 | 4.78 | Control | III-IV |
| 20 | M | 84 | 10 | Control | III-IV |
| 21 | M | 84 | NA | Control | I-II |
| 22 | M | 85 | 21 | Control | III-IV |
| 23 | F | 89 | 3.41 | Control | III-IV |
| 24 | M | 81 | 7.05 | Control | III-IV |
| 25 | F | 85 | NA | Control | I-II |
| 26 | F | 80 | 5.5 | Control | I-II |
| 27 | M | 95 | 3 | Control | III-IV |
| 28 | M | 80 | 1.91 | Control | I-II |
| 29 | F | 79 | 3 | Control | III-IV |
| 30 | M | 66 | 26.92 | Control | I-II |
| 31 | F | 97 | NA | Control | III-IV |
| 32 | M | 95 | 0 | Control | III-IV |
| 33 | F | 93 | NA | Control | I-II |
| 34 | M | 84 | NA | Control | III-IV |
| 35 | M | 86 | 2.71 | ADNC | V-VI |
| 36 | M | 83 | 3.75 | ADNC | V-VI |
| 37 | F | 92 | 3.16 | ADNC | V-VI |
| 38 | M | 84 | 4.4 | ADNC | V-VI |
| 39 | M | 93 | NA | ADNC | III-IV |
| 40 | F | 91 | 13.81 | ADNC | V-VI |
| 41 | M | 86 | 2.46 | ADNC | III-IV |
| 42 | F | 96 | 22.25 | ADNC | V-VI |
| 43 | F | 90 | 5.5 | ADNC | V-VI |
| 44 | F | 90 | 13.08 | ADNC | III-IV |
| 45 | M | 98 | 2.4 | ADNC | III-IV |
| 46 | F | 79 | 19.83 | ADNC | V-VI |
| 47 | M | 86 | 3 | ADNC | V-VI |
| 48 | F | 80 | 6 | ADNC | V-VI |
| 49 | M | 72 | NA | ADNC | V-VI |
| 50 | M | 71 | 4.41 | ADNC | V-VI |
| 51 | F | 102 | NA | ADNC | V-VI |
| 52 | F | 89 | NA | ADNC | V-VI |
| 53 | F | 89 | 2.66 | ADNC | V-VI |
| 54 | F | 94 | 5.5 | ADNC | III-IV |
| 55 | M | 85 | 17.75 | PART | III-IV |
| 56 | F | 93 | 2 | PART | III-IV |
| 57 | F | 103 | 4.4 | PART | IV-V |
| 58 | M | 75 | 4.16 | PSP | N/A |
| 59 | F | 68 | 5 | PSP | N/A |
| 60 | M | 70 | 6.91 | PSP | N/A |
| 61 | M | 70 | 0.66 | PSP | N/A |
| 62 | M | 55 | 1.5 | PSP | N/A |
| 63 | M | 67 | 1 | PSP | N/A |
| 64 | M | 69 | 0 | PSP | N/A |
| 65 | M | 71 | 0 | PSP | N/A |

F = Female; M = Male; yr = year; PMI = Postmortem Interval; hr = hour; NA = Not available; NPDX = Neuropathological diagnosis; ADNC = Alzheimer's Disease Neuropathological Changes ; PART = Primary Aged Related Tauopathy; PSP = Progressive Supranuclear Palsy; N/A = Not applicable.

**Supplementary Table 2. Antibodies used in this study**

| Antibody | Host | Manufacturer | Catalog # | IHC Dilution | IF Dilution | WB Dilution |
|---|---|---|---|---|---|---|
| NSun2 | Rabbit | Proteintech | 20854-I-AP | 1:400 | 1:100 | 1:1000 |
| NSun2 | Rabbit | ThermoFisher Scientific | 702036 | - | 1:250 | 1:500 |
| NSun2 | Mouse | Proteintech | 66580-I-Ig | - | 1:500 | - |
| Mettl3 | Rat | Abcam | ab195352 | - | - | 1:1000 |
| m6A | Mouse | Synpatic systems | 202 111 | - | - | - |
| beta-actin | Mouse | Sigma | A1978 | - | - | 1:2000 |
| beta-III-tubulin | Rabbit | Millipore | 04-1049 | - | - | 1:2000 |
| Pan-tau | Rabbit | Agilent-Dako | A0024 | - | - | 1:2000 |
| Phospho-tau (Thr181) | Mouse | ThermoFisher Scientific | MN1050 | - | - | 1:500 |
| Phospho-tau (Ser202, Thr205) (AT8) | Rabbit | ThermoFisher Scientific | MN1020 | | 1:200 | 1:500 |
| Phospho-tau (Ser214) | Rabbit | ThermoFisher Scientific | 44-742G | - | 1:50 | 1:500 |
| Phospho-tau (Thr231) | Rabbit | ThermoFisher Scientific | 701056 | - | - | 1:500 |
| Phospho-tau (Thr262) | Rabbit | ThermoFisher Scientific | 44-750G | - | - | 1:500 |
| Phospho-tau (Ser396, Ser404) (PHF1) | Mouse | Gift from Peter Davies | - | - | - | 1:1000 |
| Phospho-tau (Ser 199, Ser 202) | Rabbit | ThermoFisher Scientific | 44-768G | - | - | 1:500 |
| MAP2 | Chicken | Abcam | ab92434 | - | 1:500 | - |
| APP (6E10) | Mouse | Biolegend | SIG-39320 | - | - | 1:2500 |
| Anti-Rabbit HRP | Rabbit | Kindle Biosciences | R1006 | - | - | 1:1000 |
| Anti-Mouse HRP | Mouse | Kindle Biosciences | R1005 | - | - | 1:1000 |
| Alexa Fluor 568 goat anti-rabbit IgG | Rabbit | Thermofisher | A11011 | - | 1:1000 | - |
| Alexa Fluor 488 goat anti-mouse IgG | Mouse | Thermofisher | A11001 | - | 1:1000 | - |
| Alexa Fluor 647 goat anti-chicken IgG | Chicken | Thermofisher | A21449 | - | 1:1000 | - |

IHC = immunohistochemistry; IF = immunofluorescence; WB = Western Blot
